# Abalone genomics reveals an ancient asymmetry axis and the multifunctionality of the mantle

**DOI:** 10.64898/2026.01.03.697505

**Authors:** Xiaotong Wang, Chunhui Li, Peidong Xin, Meiwei Zhang, Baoyu Huang, Tiantian Chen, Siyi Li, Yijing Han, Lei Wei, Xuekai Zhang, Yaqiong Liu, Chenglong Zhu, Wenjie Xu, Zhiyong Yue, Wenshi Xue, Wenchao Yu, Peng Zhang, Lingling Li, Mengqiang Yuan, Deyang Tian, Jiyan Qi, Qiang Qiu, Kun Wang, Ximing Guo, Chenguang Feng

## Abstract

Gastropod diversification represents one of the most spectacular evolutionary radiations, underpinned by a fundamentally asymmetric body plan and diverse pigmentation patterns. To understand the genetic underpinnings of these cardinal traits, we constructed a chromosome-level genome assembly for the rainbow abalone, *Haliotis iris*. Integrating systematic genomics, histoembryology, and molecular assays, we uncover an ancient regulatory axis, conserved for over 544 million years, in which the long non-coding RNA *lncRNA1* functions as a competing endogenous RNA to shield *pitx* transcripts from miRNA-mediated degradation, thereby driving the establishment of asymmetric body patterning, including the mantle. Furthermore, we identify a *wnt-mitf-tyr* cascade as the driver of melanogenesis and demonstrate that the mantle is the primary site of melanin synthesis. Concurrently, we reveal the mantle as a multifunctional organ possessing mechanosensory capabilities. These findings elucidate the ancient genetic toolkit governing trait evolution in gastropods.

**Teaser:** A deeply conserved RNA switch directs gastropod body plans and reveals the mantle’s hidden talents.

## Introduction

Gastropods, with their unparalleled radiation into over 80,000 extant species, represent a pinnacle case of morphological and ecological diversification within the Mollusca ^1,2^. Their hallmark is an asymmetric body plan forged by torsion and subsequent shell coiling, a developmental process that reorients the visceral mass relative to the head and foot, fundamentally shaping their morphology and ecological adaptations ^3,4^. This asymmetry, along with melanin-based pigmentation critical for photoprotection, camouflage, and signaling, underpins their phenotypic diversity ^5–7^. These features, asymmetry and pigmentation, are not merely curiosities but windows into the genetic and molecular mechanisms that have driven the diversification of the gastropod clade.

Despite significant advances in molluscan genomics ^8^, key gaps remain in our understanding of these core gastropod traits. The molecular basis of organ asymmetry remains elusive; while *pitx*, known to determine left-right asymmetry in bilaterians ^9,10^, has been reported to be associated with coiling in gastropods ^4,11,12^, its regulatory architecture in gastropods remains inadequately comprehended. Similarly, melanin synthesis, well-described in vertebrates and a few invertebrates through the gene *tyr* ^13–16^, is understudied in gastropods, leaving its pathway conservation and regulation unclear ^17,18^.

The rainbow abalone, *Haliotis iris*, offers a unique system for studying gastropod asymmetry and melanogenesis. A hallmark of the Haliotidae is a partial, ∼90-degree detorsion that follows the canonical 180-degree gastropod torsion ^19,20^. This process directly results in an anterior mantle cleft with asymmetric lobes (right > left), a defining morphological synapomorphy that differs from most gastropods with a unified mantle ^20,21^. Complementing this morphological novelty is its suitability for investigating melanogenesis. The species exhibits a striking eumelanin-based black phenotype (Fig. 1A) ^22^, yet includes the natural variation of unpigmented regions in some individuals (see below), providing a tractable system for genetic dissection. As an ecologically significant species in New Zealand’s intertidal zones (Fig. 1A) and one of the top three abalone species in global trade ^23–25^, *H. iris* combines profound evolutionary interest with substantial economic and cultural relevance.

**Fig. 1.**
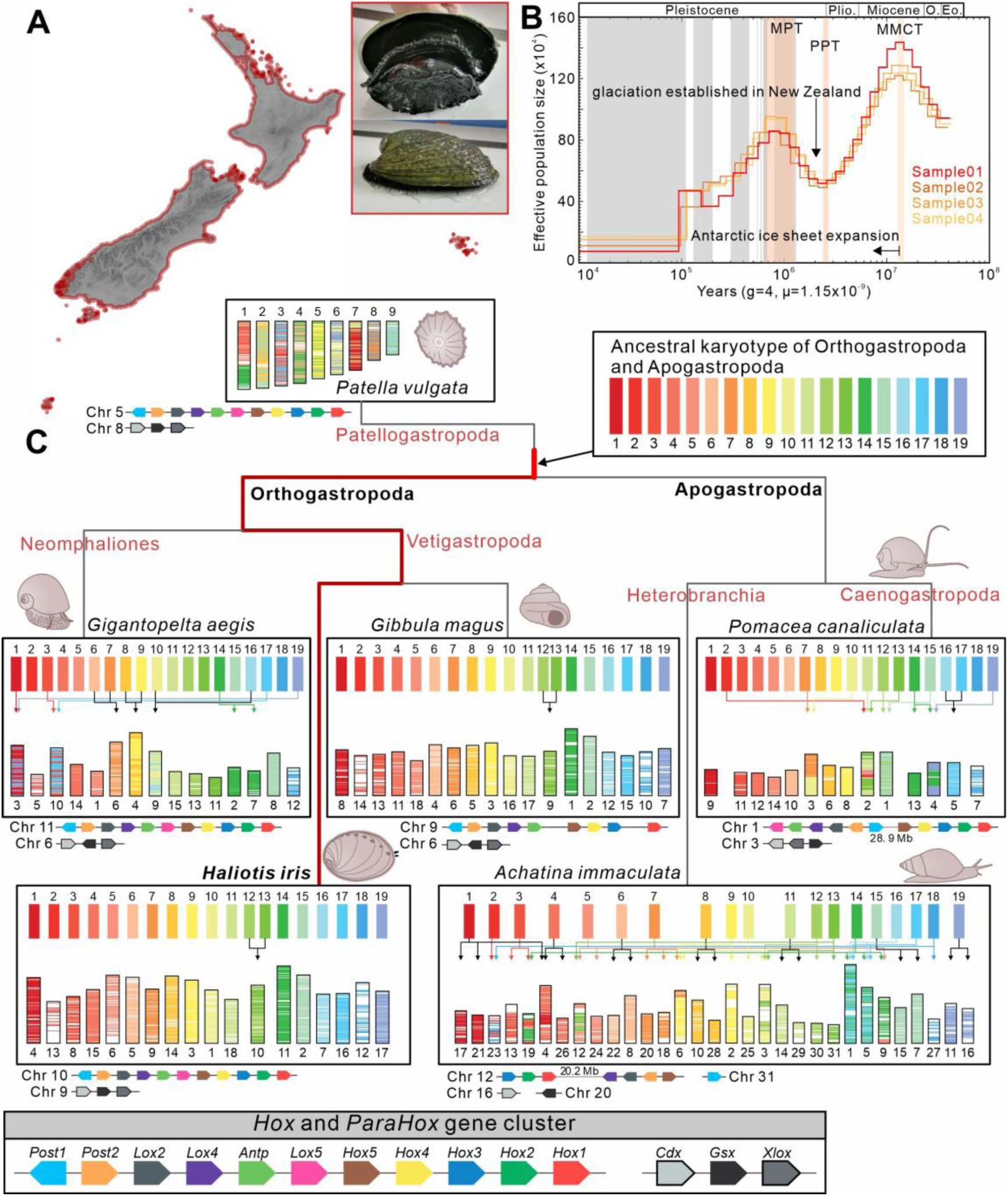
Ecological distribution and evolutionary analyses of rainbow abalone *H. iris*. (A) Morphology and geographical distribution. (B) Historical effective population size; four colored lines represent the demographic history reflected by four individual *H. iris*; orange bars indicate major paleoclimatic events with annotations; gray bars represent Pleistocene glacial periods. (C) Distribution of ancestral proto-chromosomal fragments and chromosomal organization of *Hox* and *ParaHox* genes in the genomes of six representative gastropod species.

Leveraging a high-quality, chromosome-level genome assembly of *H. iris* and comprehensive comparative analyses across gastropods, we delineated genomic features consistent with its evolutionary conservatism. We subsequently dissected the molecular underpinnings of two pronounced features of gastropods, developmental asymmetry and pigmentation. As these cardinal traits are both linked to the mantle, our investigation further uncovered molecular evidence for its multifunctionality. Finally, this work also revealed a severe demographic bottleneck in *H. iris* over the last 110,000 years, resulting in a small effective population size and highlighting the imperative for conservation of this ecologically and economically important species.

## Results and Discussion

### Genome information

A chromosome-level *H. iris* genome assembly was obtained using a combination of Oxford Nanopore (∼148.6x coverage), Illumina (∼178.3x coverage), and Hi-C (∼186.4x coverage) sequencing reads (Supplementary Table S1). The final assembly spanned 1,150.57 Mb, with a scaffold N50 of 66.58 Mb and a contig N50 of 12.77 Mb (Table 1). A total of 99.1% of the assembled bases were anchored to 18 pseudo-chromosomes (Supplementary Fig. S1), consistent with the expected haploid chromosome number and genome size ^26,27^ (Supplementary Fig. S2). The BUSCO assessment indicated high completeness and accuracy, with 96.8% of metazoan conserved core genes covered (Supplementary Table S2) and a single-base error rate of less than 0.0627 per megabase (Table 1). Genome annotation identified 20,994 protein-coding genes and 566.94 Mb of repetitive sequences, with the latter accounting for 49.3% of the genome (Table 1; Supplementary Table S3). The protein-coding genes included 98.7% of metazoan BUSCOs (Supplementary Table S2), and 91.18% of these genes were supported by transcript evidence (17,626 genes) or homology in public databases (18,095 genes) (Supplementary Table S4). The quality of the *H. iris* genome surpasses that of most published molluscan genomes and represents the highest-quality genome for the Haliotidae family to date (Supplementary Fig. S3), providing a robust foundation for subsequent analyses.

**Table 1.** Assembly statistics of the rainbow abalone genome.

| Feature | Statistics |
| --- | --- |
| Assembly size (Mb) | 1,150.57 |
| Scaffold N50 (Mb) | 66.58 |
| Contig N50 (Mb) | 12.77 |
| Longest scaffold (Mb) | 84.81 |
| Longest contig (Mb) | 40.58 |
| Haploid number of *pseudo*-chromosomes | 18 |
| Chromosome bases/total length (%) | 99.09 |
| Base error rate in assembly (per Mb) | 0.0627 |
| Repeats (Mb) | 566.94 |
| Protein-coding genes | 20,994 |
| BUSCO of assembly (%) | 96.8 |
| BUSCO of Protein-coding genes (%) | 98.7 |

### Demographic history and chromosome evolution

Phylogenetic analysis revealed that *H. iris* diverged from its sibling species, *H. rufescens*, around 39.9 Ma (95% CI: 23.19-56.36 Ma; Supplementary Fig. S4). We reconstructed the demographic history of rainbow abalone over the past 40 Ma by estimating effective population size (Fig. 1B). Over this period, the population experienced three major fluctuations and one severe bottleneck. It declined from its peak during the Middle Miocene Climate Transition (MMCT) ^28,29^ around 14.55-12.16 Ma; rebounded after the warmer Plio-Pleistocene Transition (PPT) ^30^ from 2.65-1.96 Ma; dropped again after the cooler Mid-Pleistocene Transition (MPT) ^31^ around 0.93-0.71 Ma; and underwent a severe contraction from ∼0.11 Ma, coinciding with the onset of the Würm glaciation ^32^, which resulted in a severe bottleneck with a small effective population size of 7.47-16.89 (x10⁴). Re-sequencing of three additional rainbow abalone individuals validated these demographic dynamics (Fig. 1B).

Chromosome evolution was investigated by comparing the karyotypes of six representative gastropod species: *Patella vulgata*, *Gigantopelta aegis*, *H. iris*, *Gibbula magus*, *Achatina immaculata*, and *Pomacea canaliculate* (Fig. 1C). Reconstruction of the ancestral karyotype for Orthogastropoda and Apogastropoda (2n = 38; Supplementary Table S5) revealed that while chromosomal fusions and fissions are frequent across gastropods, the Vetigastropoda lineage, which includes *H. iris*, exhibits remarkable karyotypic conservation with only one inferred proto-chromosome fusion event. Furthermore, we examined the arrangement of *Hox* and *ParaHox* gene clusters, finding that *H. iris* has retained the conserved ancestral configuration for both (Fig. 1C). Taken together, the abalone preserves some ancient genetic features, representing an archetypal, evolutionarily conserved lineage of gastropods.

An intriguing correlation emerges wherein lineages with few karyotypic changes (e.g., *H. iris* and *G. magus*) are typically open-water broadcast spawners capable of wide larval dispersal. In contrast, lineages with extensive karyotypic rearrangements (e.g., *A. aegis*, *A. immaculata*, and *P. canaliculata*) are often land snails or deep-sea hydrothermal vent species with limited larval dispersal. One explanation is that limited larval dispersal increases the probability of inbreeding, thereby enhancing the fixation rate of chromosomal mutations. This inbreeding hypothesis is consistent with observations in Pectinidae scallops ^33^. Further supporting this, the scallop *Patinopecten yessoensis*, a broadcast spawner with wide larval dispersal, possesses a highly conserved karyotype similar to the ancestral bilaterian karyotype, which aligns with the prediction that reduced inbreeding constrains karyotypic change ^34^. Further studies are needed to fully understand the mechanisms and evolutionary consequences of karyotypic evolution.

### A lncRNA-*pitx* axis governs mantle asymmetry

The abalone mantle is a classic example of organ asymmetry in a bilaterally symmetric animal, with the right side being larger than the left (Fig. 2A). To identify the genetic determinants of this asymmetry, we compared the transcriptomes of paired organs: the asymmetric left and right mantle lobes and the symmetric left and right cephalic tentacles. No genes were differentially expressed between the symmetric cephalic tentacles (Fig. 2A; Supplementary Table S6). In contrast, a single protein-coding gene, *pitx*, was differentially expressed between the asymmetric mantle lobes; its expression was high in the smaller left lobe but nearly undetectable in the larger right lobe (Fig. 2A; Supplementary Fig. S5; Supplementary Table S7). This difference in expression pattern was also observed in a sibling species, *H. discus* (Fig. 2B). Across several bivalve species, *pitx* similarly showed higher expression on the smaller side of the mantle, except for *Scapharca broughtonii*, where the mantle lobes are of comparable size (Fig. 2B). These findings implicate *pitx* as a key transcription factor regulating molluscan mantle symmetry *via* inhibitory effects on tissue growth.

**Fig. 2.**
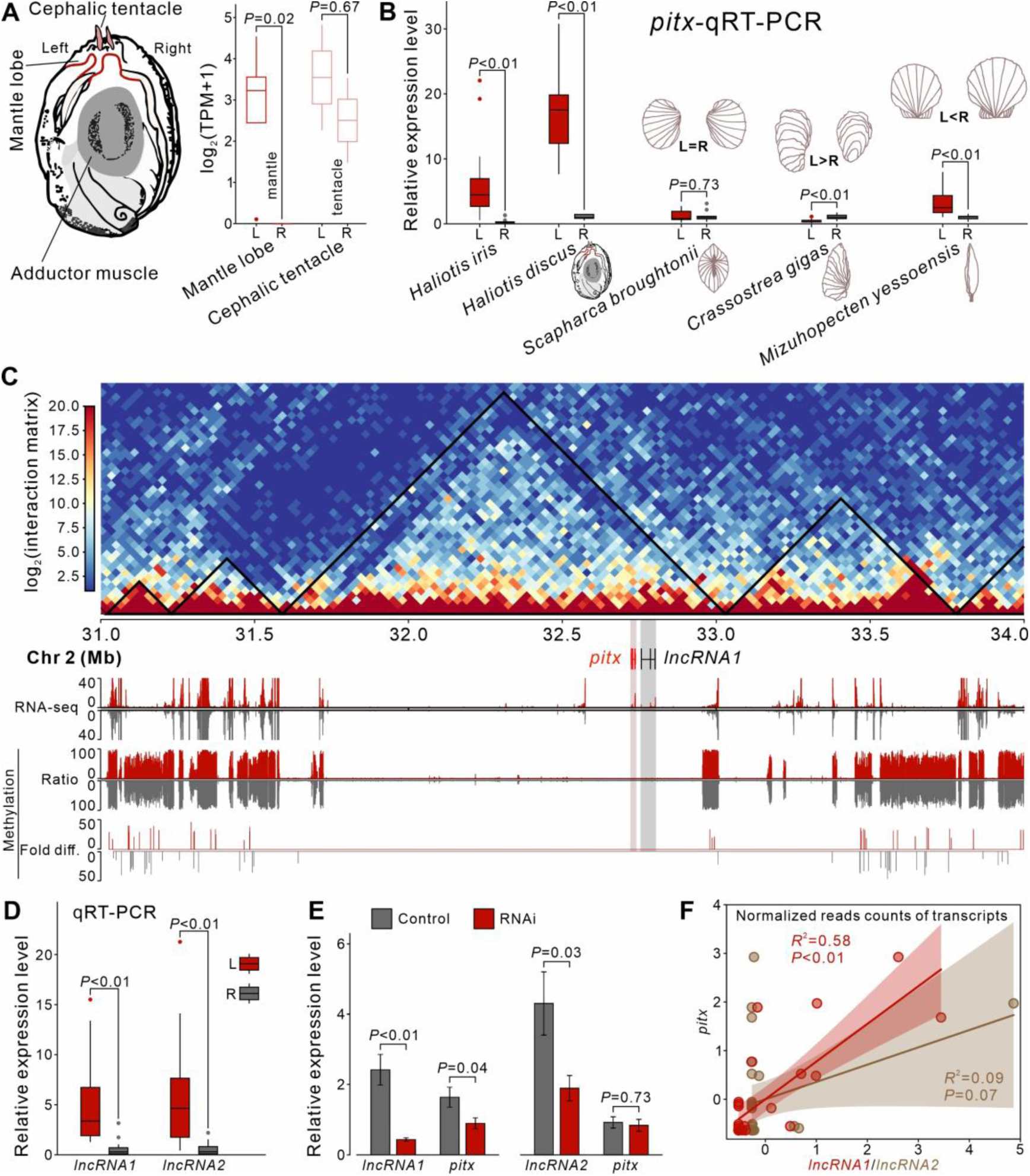
Asymmetric expression of *pitx* and lncRNAs in mantle lobe pairs. (A) Dorsal view schematic of the shell-less abalone and *pitx* expression differences between left and right mantle lobes and cephalic tentacles. (B) qRT-PCR comparing *pitx* expression in left *vs* right mantle lobes of four other mollusks. (C) Chromatin interactions, transcripts, and methylation in the Chr2:31-34 Mb region surrounding *pitx*; black lines in interaction plot indicate TADs; *pitx* and *lncRNA1* models shown in red and black, respectively; RNA-seq and methylation tracks show left side in red (top half) and right side in gray (bottom half). (D) qRT-PCR comparing the expression of two lncRNAs in left *vs* right mantle lobes of rainbow abalone. (E) qRT-PCR results showing that only *lncRNA1* knockdown reduces *pitx* expression. (F) General linear regression across the expression of lncRNAs *vs. pitx* in tissues of rainbow abalone.

Given that *pitx* was the only differentially expressed protein-coding gene, we sought to understand the regulation of its asymmetric expression. In oysters, we previously found that asymmetric *pitx* expression correlated with differential DNA methylation in its upstream regulatory region ^35^. In *H. iris*, however, we observed extremely low methylation levels across a 150 kb region flanking *pitx* with no difference between the left and right sides (Fig. 2C; Supplementary Fig. S5), suggesting a novel regulatory mechanism is at play. A subsequent screen for non-coding RNAs (ncRNAs) identified two previously unannotated long ncRNAs (lncRNAs) that were differentially expressed between the left and right mantle lobes (Supplementary Table S8). One of these, *lncRNA1*, is located 18.6 kb downstream of *pitx* and mirrors its expression pattern, being highly expressed in the smaller left lobe (Fig. 2C, 2D; Supplementary Tables S8, S9). The other, *lncRNA2*, resides on a different chromosome (Chr 12). Neither lncRNA showed differential expression in the symmetrical cephalic tentacles (Supplementary Table S8), further implicating them in the specific regulation of mantle asymmetry. While both lncRNAs showed elevated expression in the left mantle lobe, similar to *pitx* (Fig. 2D), only the knockdown of *lncRNA1* perturbed *pitx* expression levels, revealing a positive correlation (Fig. 2E; Supplementary Table S10). This correlation was observed across multiple tissues (Fig. 2F), indicating that *lncRNA1* interacts with *pitx* in regulating tissue growth and mantle asymmetry in abalone.

### An ancient *lncRNA1*-miRNAs-*pitx* cascade controls gastropod asymmetry

To dissect the molecular mechanism of the *lncRNA1-pitx* axis, we performed a histological survey of the left anterior mantle lobe in *H. iris* (Fig. 3A). *In situ* hybridization revealed that both *pitx* and *lncRNA1* are exclusively expressed in the mantle fold region and are highly co-localized in the cytoplasm (Fig. 3B, 3C). The cytoplasmic co-localization suggested that *lncRNA1* might function as a “guardian” for *pitx* mRNA, consistent with the classic competing endogenous RNA (ceRNA) model, where lncRNAs sequester microRNAs (miRNAs) ^36,37^. To test this hypothesis, we used HyPro-seq ^38^ with an *lncRNA1*-specific probe to capture potentially interacting miRNAs (Supplementary Tables S11, S12). We identified three high-confidence miRNAs that were also localized to the mantle fold region (Fig. 3D, 3E; Supplementary Fig. S6). A dual-luciferase reporter assay confirmed that all three miRNAs could bind to both *pitx* and *lncRNA1* transcripts and reduce their stability (Fig. 3F; Supplementary Table S13). This supports a model where *lncRNA1* acts as a molecular sponge, sequestering these miRNAs to protect *pitx* mRNA from degradation (Fig. 3G). Concurrently, analysis of chromatin interactions revealed a strong loop signal in the right mantle between *lncRNA1* and a 707.5 kb downstream TAD boundary, a structure absent in the left mantle (Fig. 3H). As TAD boundaries are often enriched with repressive boundary proteins ^39,40^, we speculate that this differential chromatin looping may also contribute to the asymmetric expression of *pitx* and *lncRNA1*.

**Fig. 3.**
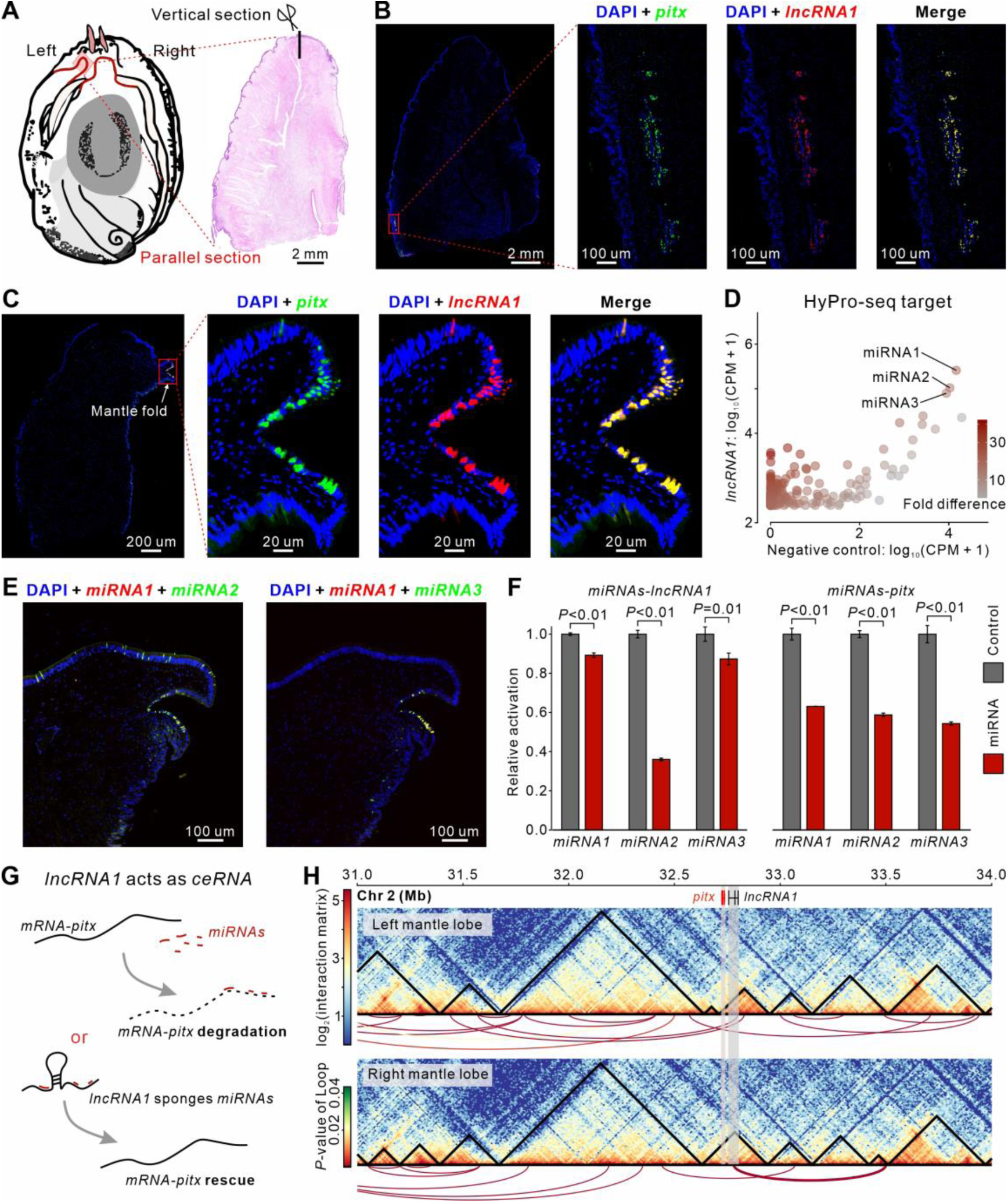
Spatial localization and interaction mechanism between *pitx* and *lncRNA1*. (A) Position diagram and parallel HE section of the left mantle lobe. (B and C) *In situ* hybridization results for the *pitx* and *lncRNA1* genes in the parallel (B) and vertical (C) sections of the left mantle lobe. (D) The CPM abundance of miRNAs captured by the HyPro-seq assay using the *lncRNA1* (y-axis) and random sequence (x-axis) probes. (E) *In situ* hybridization results for the three miRNAs in the vertical sections of the left mantle lobe. (F) The regulatory effect of miRNAs on *lncRNA1* and *pitx* gene was measured using the dual-luciferase reporter assay system. (G) Schematic diagram of the *lncRNA1*-miRNAs-*pitx* axis. (H) Chromatin interactions in the Chr2:31-34 Mb region between left and right mantle lobes; black lines in the interaction plot indicate TADs; *pitx* and *lncRNA1* gene models shown in red and black, respectively; lines below the heatmap represent the DNA loops between HIC units, and the color of the line tells the corresponding *p*-value.

These findings provide the first evidence for a *lncRNA1*-miRNAs-*pitx* axis regulating tissue growth and mantle asymmetry in abalone. Transcriptome analysis showed that both *lncRNA1* and *pitx* expression initiates at the gastrula stage (Fig. 4A), and they remain highly co-localized throughout development, closely associated with the formation of the mantle and other organs (Fig. 4B). This indicates the *lncRNA1*-miRNAs-*pitx* axis is deeply integrated into the establishment of the abalone body plan. On a macroevolutionary scale, ncRNAs are often poorly conserved, making cross-lineage comparisons difficult ^41,42^. However, our analysis of gene synteny revealed that the *lncRNA1* and *pitx* loci are highly linked throughout gastropod evolution (except in Apogastropoda) and are consistently located on the ancestral chromosome 15 of Orthogastropoda and Apogastropoda and their descendants (Fig. 4C; Supplementary Table S14). This remarkable synteny suggests that the *lncRNA1-*miRNA*-pitx* axis is an ancient and highly conserved regulatory mechanism for body patterning in gastropods, dating back 544.4 million years (Supplementary Fig. S4).

**Fig. 4.**
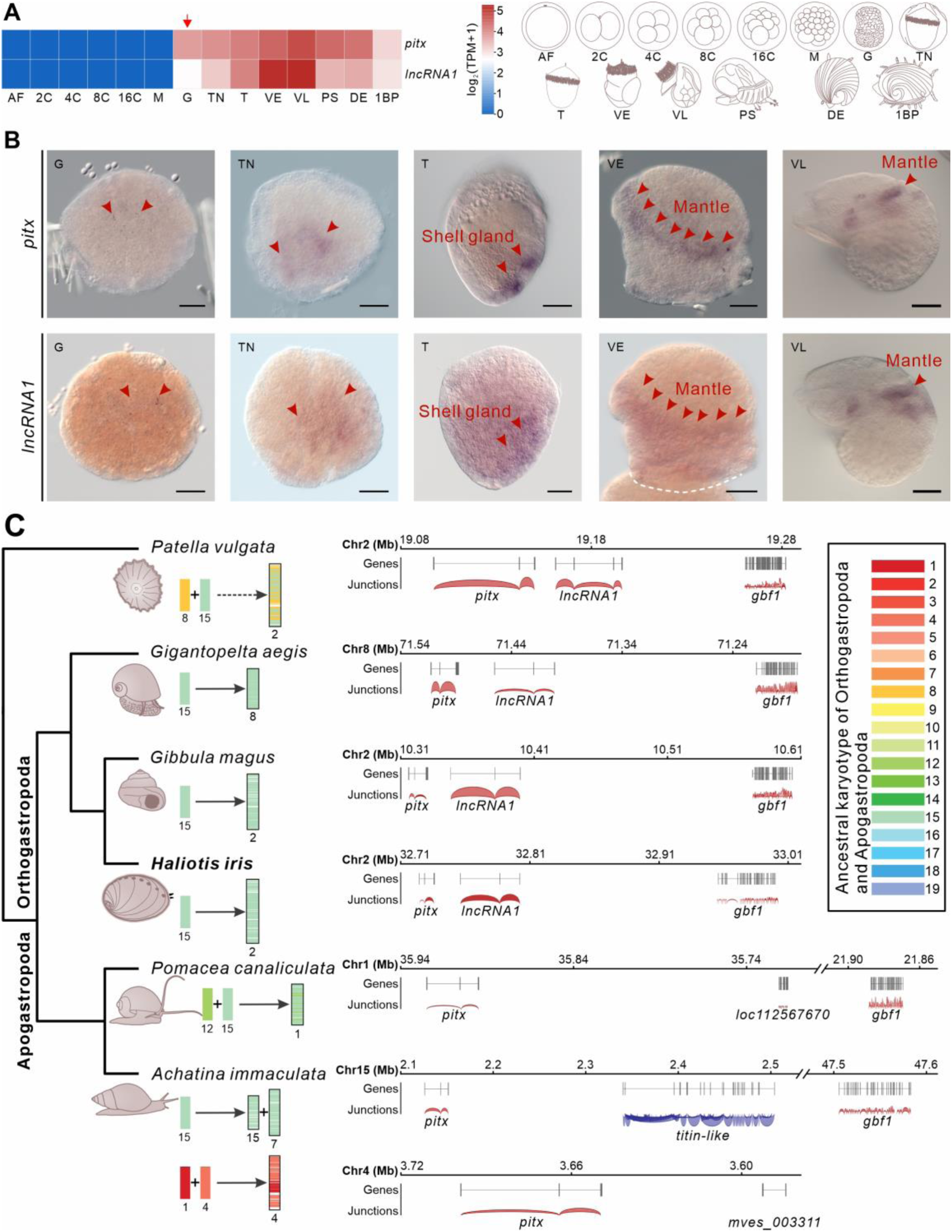
An ancient *lncRNA1-pitx* axis regulates the body plan of gastropods. (A) Expression dynamics of *pitx* and *lncRNA1* during *Haliotis discus* embryonic development; AF: artificial fertilization; 2C: 2-cell; 4C: 4-cell; 8C: 8-cell; 16C: 16-cell; M: morula; G: gastrula; TN: non-hatched trochophore; T: trochophore; VE: early veliger; VL: late veliger; PS: peristomial shell larva; DE: differentiation epipodes; 1BP: emergence of 1st breathing pore. (B) *In situ* hybridization showed that *pitx* and *lncRNA1* had consistent expression sites across embryonic stages. (C) The conserved synteny between *pitx* and *lncRNA1* can be traced back to the ancestor of gastropods; their loci are consistently located on the ancestral chromosome 15 of Orthogastropoda and Apogastropoda and its descendants; blue junctions indicate that the genes are located on complementary strands.

### Chitin-binding proteins are linked to melanin deposition

Melanin is a ubiquitous pigment in metazoans, responsible for the coloration of skin, hair, and eyes, and serving functions in photoprotection and immunity ^5,7,43,44^. While melanin synthesis in vertebrates proceeds via a well-characterized tyrosine metabolic pathway, the pathway in most other taxa remains poorly understood ^13–15^. The all-black skin of *H. iris* distinguishes it from other abalone species (Fig. 1A); however, some individuals exhibit an unpigmented lower pedal region (Fig. 5A). Electron microscopy confirmed a lack of melanin in this unpigmented tissue (Supplementary Fig. S7). This natural variation in melanin deposition offers an excellent model for investigating melanogenesis in gastropods. We performed comparative transcriptomics between pigmented and unpigmented tissues from both normal and variant individuals, comparing pigmented epithelium to unpigmented muscle and pigmented epithelium to unpigmented epithelium (Fig. 5A). Results showed 1,840 up-and 1,055 down-regulated genes in pigmented regions in either comparison (Supplementary Tables S15, S16). However, no known melanogenic genes were identified among them, and the key gene *tyr* was undetectable in any transcripts from these tissues (Supplementary Table S17). This suggests that melanin in the lower pedal is not synthesized *in situ* and that the color variant is likely determined by factors affecting melanin deposition rather than synthesis.

**Fig. 5.**
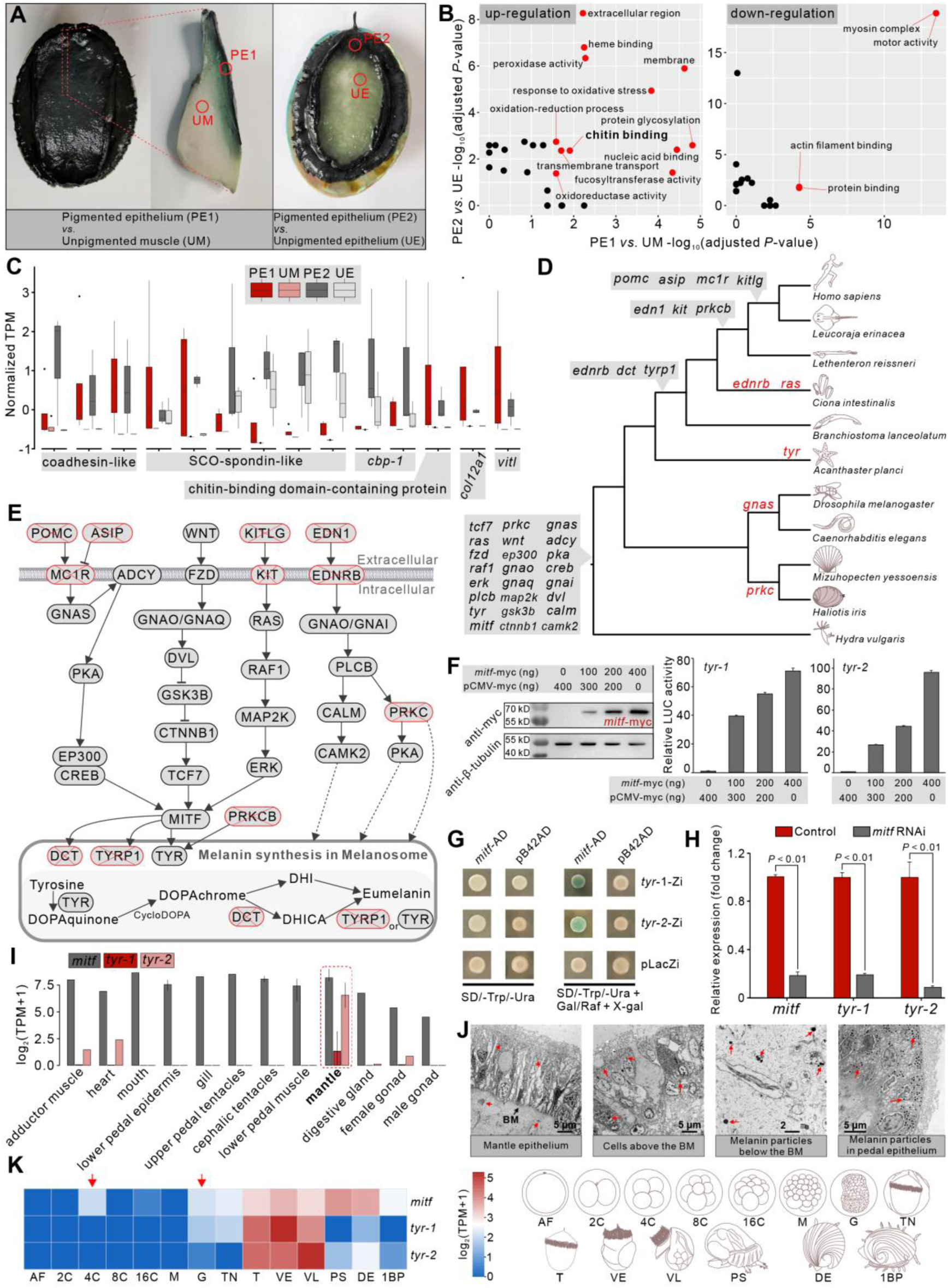
Analyses of melanin-related genes in rainbow abalone. (A) Illustration of normal and pigment-defective rainbow abalone variants and sampled tissues. (B) GO enrichment of genes differentially expressed in pigmented *vs* unpigmented tissues; red dots indicate terms significant in both comparisons, black dots indicate terms only significant in one comparison. (C) Expression levels of chitin-binding related genes across four tissues. (D) Origins (gray background) and losses (red) of melanogenic genes across metazoan phylogeny. (E) Schematic of known eumelanin synthesis and regulation; genes lost in rainbow abalone are marked with red crosses. (F) Dual luciferase assays showing transcriptional regulation of *tyr* genes by rainbow abalone *mitf*; Western blots on the left show increased MITF expression with higher *mitf*-myc vector concentrations. (G) Yeast one-hybrid assays validating *mitf* regulation of *tyr*; green patches indicate positive results. (H) qRT-PCR results showing *mitf* knockdown reduces *tyr* expression. (I) *mitf* and *tyr* expression across rainbow abalone tissues. (J) Transmission electron micrographs of abalone mantle and foot; red arrows indicate melanin particles, BM: basement membrane. (K) Expression dynamics of *mitf* and *tyr* during *Haliotis discus* embryonic development; embryonic stages indicated by abbreviations refer to Fig. 4A.

Transcriptome analyses did, however, reveal an up-regulation of chitin-binding genes, such as *coadhesin-like* and *SCO-spondin-like*, in all melanin-deposited tissues (Fig. 5B-5C). These genes were also enriched in GO terms associated with the cell membrane and extracellular region. Previous studies have found melanin deposited within cross-linked chitin structures in cuttlefish beaks, oyster shells, and silkworm epidermis ^45–48^. This implicates the involvement of chitin-rich matrices in melanin localization across invertebrates.

### Identification of the core melanogenesis pathway

Since transcriptomic comparisons of pigmented tissues were uninformative for identifying melanogenic genes, we used an evolutionary genomics approach to uncover the regulatory cascade. Melanogenesis mediated by *tyr* is evolutionarily conserved across metazoans ^43,44,49^, implying an ancient regulatory pathway. Melanin in *H. iris* is eumelanin, which is also *tyr*-driven in mollusks ^22,49,50^. We therefore investigated the origin and loss of all genes in the eumelanogenic pathway (ko04916) across metazoans (Fig. 5D). Of the three canonical eumelanogenic genes (*dct*, *tyrp1*, and *tyr*), only *tyr* exists in non-chordate metazoans. This suggests that the synthesis of melanin in *H. iris* is dominated by *tyr* and proceeds through the 5,6-dihydroxyindole (DHI) approach (Fig. 5E).

The *H. iris* genome contains two *tyr* paralogs (*tyr-1* and *tyr-2*). Investigating upstream regulators, we found that only the transcription factor *mitf* is conserved across non-chordate metazoans (Fig. 5D-5E). To test whether *mitf* regulates *tyr* in *H. iris*, we performed dual-luciferase reporter assays in human embryonic kidney 293T (HEK293T) cells (Supplementary Fig. S8). The results showed that *mitf* positively regulated both *tyr* paralogs (Fig. 5F; Supplementary Fig. S9; Supplementary Table S18). This *mitf-tyr* interaction was further validated by yeast one-hybrid assays (Fig. 5G). Furthermore, knockdown of *mitf* led to a significant downregulation of both *tyr-1* and *tyr-2* (Fig. 5H; Supplementary Table S19). We therefore conclude that *mitf* is the key transcriptional activator of *tyr* in *H. iris*. Regarding the upstream regulation of *mitf*, our genomic survey revealed that key components of pathways other than the *wnt* signaling cascade are absent in *H. iris* (Fig. 5E). In contrast, all core *wnt* pathway genes are conserved across metazoan lineages (Fig. 5D), suggesting that the *wnt-mitf-tyr* cascade may represent an ancient, ancestral pathway for melanogenesis in metazoans.

### The mantle is the primary site of melanogenesis

If melanin in the lower pedal is not synthesized locally, what is its origin? To identify the source tissue, we examined the expression patterns of the key melanogenic genes, *mitf* and *tyr*, across different organs. The two *tyr* paralogs were predominantly expressed in the mantle and were barely detectable in most other organs, including the lower pedal (Fig. 5I). Although *tyr-2* showed some expression in the heart, adductor muscle, and female gonad, there is no evidence that connective tissues link these organs to the lower pedal skin. This strongly implies that cutaneous melanin in *H. iris* is primarily produced in the mantle and subsequently transported. Electron microscopy confirmed the presence of noticeable melanin particles beneath the basement membrane of the mantle epithelium (Fig. 5J), validating the melanogenic function of the mantle.

Intriguingly, while the mantle is the primary site of *tyr* expression in adults (especially *tyr-1*, Fig. 5I), *tyr* expression begins at the gastrula stage (Fig. 5K), which precedes the formation of the mantle at the shell gland of the trochophore larva ^51,52^. This temporal discrepancy may be attributed to either the pleiotropy of *tyr* or a deeper, yet unknown, link between *tyr* and mantle development, such as cues that the mantle forms earlier.

### Evidence for a mechanosensory function of the mantle

In addition to its role in melanogenesis, the mantle may also possess a mechanosensory function, as suggested by the specific and high expression of multiple members of the solute carrier family 26 (Fig. 6A). These membrane transporters have diverse roles mediated by anion transport ^53^. Notably, one member, *slc26a5* (known as *prestin*), mediates electromotility in the outer hair cells of the mammalian cochlea, transducing mechanical signals into electrical signals, which is the basis of hearing (Fig. 6B) ^54^. It has been reported that *slc26a5* is consistently expressed in mechanosensory ciliated cells and/or auditory organs across phyla ^55,56^, implicating a close association with mechanosensation ^57^.

**Fig. 6.**
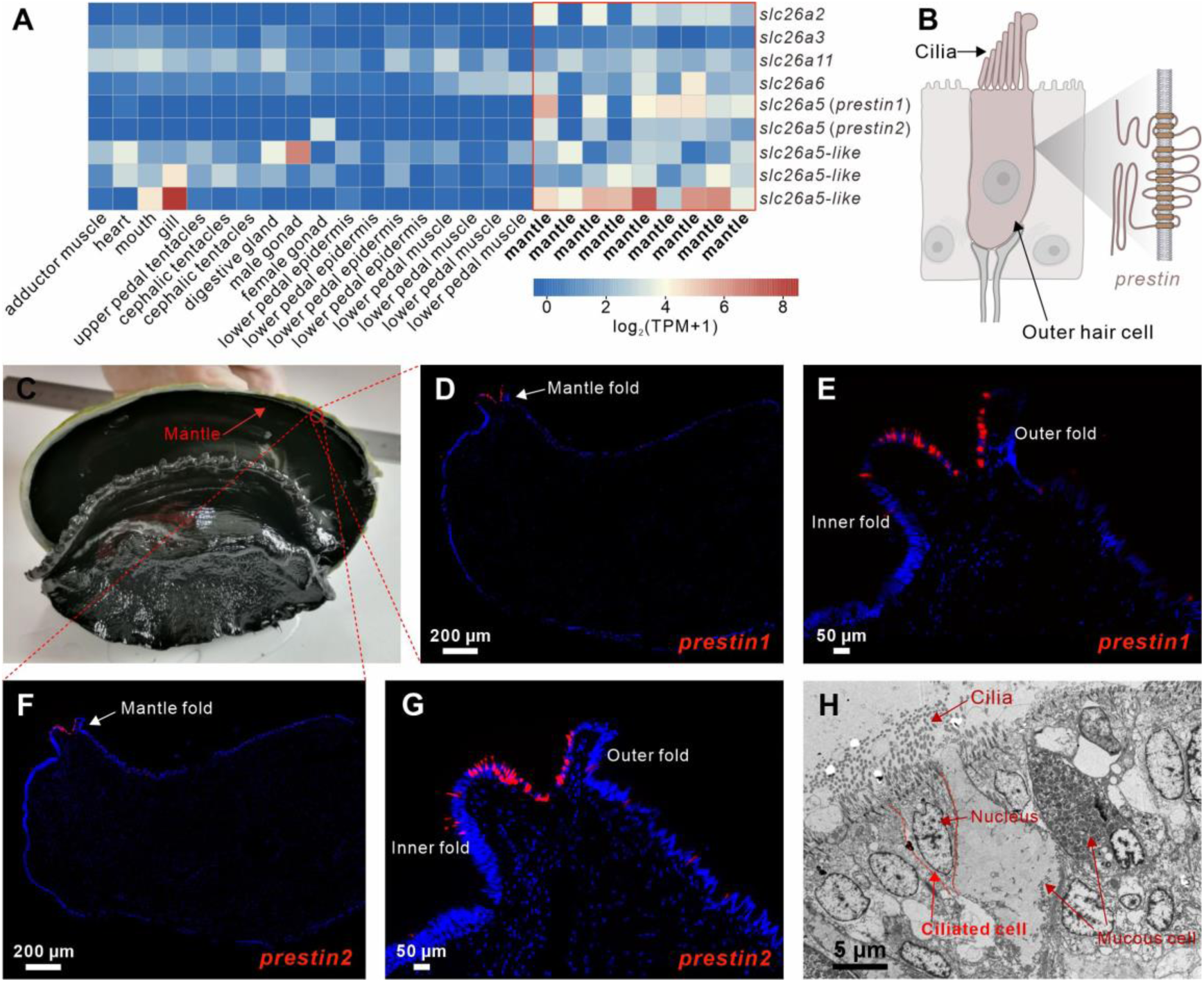
Specific expression of *slc26a5* in the rainbow abalone mantle. (A) Expression levels of solute carrier family 26 members across tissues. (B) Schematic of outer hair cell structure and *prestin* (mammalian *slc26a5*) localization in mammals. (C) Photo of the rainbow abalone mantle. (D-G) *In situ* localization of two *slc26a5* paralogs in the rainbow abalone mantle. (H) Transmission electron micrograph of the *slc26a5* expression region in the rainbow abalone mantle.

The *H. iris* genome contains two *slc26a5* genes and three *slc26a5-like* genes (Fig. 6A). In particular, *pretin1* has the same transmembrane structure as the classic *prestin* gene responsible for hearing (Supplementary Table S20) ^58^. The two *slc26a5* genes are expressed almost exclusively in the mantle, with localization specifically in the inner and outer mantle folds (Fig. 6C-6G). These folds are anatomically rich in ciliated cells (Fig. 6H). These findings suggest that the mantle may play an important role in mechanosensing in abalone.

## Conclusions

The high-quality, chromosome-level genome of the rainbow abalone, *H. iris*, opens a unique evolutionary window into gastropod evolution and diversification that have been poorly understood. Our analyses demonstrate that *H. iris* represents a genomically lineage, characterized by a stable karyotype and the retention of ancestral *Hox* and *ParaHox* gene configurations. The karyotypic conservation, which we link to its broadcast spawning life history that minimizes inbreeding and constrains chromosomal rearrangements, establishes the abalone as a pivotal model for understanding deep gastropod evolution.

A key finding of this study is the elucidation of a novel regulatory cascade governing left-right asymmetry, a fundamental process in developmental biology. We have moved beyond the established role of *pitx* to uncover a more complex, multi-layered mechanism. We demonstrate that a highly syntenic *lncRNA1* acts as a cytoplasmic ceRNA, or molecular sponge, to protect *pitx* mRNA from miRNA-mediated degradation. This *lncRNA1*-miRNA-*pitx* axis, coupled with differential chromatin looping, drives the asymmetric development of the mantles. The profound evolutionary linkage between the *pitx* and *lncRNA1* loci across gastropods suggests this regulatory module is not a recent invention but an ancient mechanism for body patterning that has been conserved for 544.4 million years.

Furthermore, our research redefines the function of the gastropod mantle, not merely an organ for shell formation, but a multifunctional organ central to both pigmentation and mechanosensation as well. We have dissected the core melanogenesis pathway in a gastropod, establishing the conserved *wnt-mitf-tyr* signaling cascade as the driver of melanin production. Crucially, we show that the mantle is the primary site of melanin synthesis, with the pigment subsequently transported to distal cutaneous tissues—a novel mechanism for skin coloration in this group. Simultaneously, the mantle exhibits highly specific expression of *slc26a5* (*prestin*) orthologs within its ciliated folds, providing evidence for a sophisticated mechanosensory function, likely for detecting water movement or vibration.

In conclusion, our comprehensive genomic and functional analyses of the rainbow abalone have not only resolved key aspects of its demographic and evolutionary history but have also uncovered deeply conserved genetic toolkits for development and physiological adaptation. These findings on organ asymmetry and the multi-function mantle advance our understanding of the evolution of complexity and innovation in one of the most diverse faunas.

## Materials and Methods

### Sample collection and sequencing

Rainbow abalone *H. iris* samples were obtained commercially and acclimated in seawater at 18°C for two days. Subsequently, the following tissues were dissected and flash-frozen in liquid nitrogen (Supplementary Table S1): upper pedal muscle (unpigmented), lower pedal epithelium (pigmented), lower pedal epithelium (unpigmented), lower pedal muscle (unpigmented), left and right mantle lobes, adductor muscle, gill, mantle, digestive gland, upper pedal tentacles, mouth, heart, male gonad, female gonad, and left and right cephalic tentacles. These samples were used for extracting genomic DNA, sequencing library construction (Oxford Nanopore long reads and Illumina 150 bp paired-end reads), transcriptomics, Hi-C, and methylome. Oxford Nanopore sequencing was performed on the PromethION platform and other libraries were sequenced on the Illumina NovaSeq 6000 platform. All sequencing was conducted at Novogene (Tianjin, China).

### Genome assembly

Illumina NGS reads were quality filtered using fastp v0.23.1 ^59^ (default parameters) and then subjected to 23-mer frequency analysis using Jellyfish v2.2.10 ^60^. GenomeScope 2.0 ^61^ was used to estimate the genome size based on the *k*-mer distribution.

Nanopore reads were quality controlled with Ontbc v1.1 (parameters: read length >1000bp, quality score >8; https://github.com/FlyPythons/ontbc) and then assembled into contigs using NextDenovo v2.4.0 with default parameters (https://github.com/Nextomics/NextDenovo). The preliminary assembly was polished in two rounds using the filtered Nanopore reads and then in three rounds using the Illumina reads with NextPolish (parameters: -min_read_len 5k -max_depth 100; https://github.com/Nextomics/NextPolish).

The Hi-C reads were aligned to the polished genome using Juicer v1.5 ^62^, and a Hi-C contact matrix was generated. The 3D-DNA v180922 ^63^ was then used to scaffold the genome into chromosome-level *pseudo*-haplotypes with default parameters. The assembly was further curated in Juicebox ^64^ to obtain the final version.

Genome completeness was assessed by benchmarking universal metazoan genes (metazoa_odb10) using BUSCO v4 ^65^. Illumina reads were aligned to the final assembly using BWA v0.7.15-r1140 ^66^. After removing ambiguous mappings, the single-base error rate was estimated using an in-house perl script.

### Transcriptome assembly and annotation

RNA-seq reads from all tissues were filtered with fastp v0.23.1 ^59^ and then assembled into transcripts using Trinity v2.8.5 ^67^. Open reading frames (ORF) were identified from the transcripts using TransDecoder v5.5.0 (https://github.com/TransDecoder/TransDecoder) with default parameters. Only transcripts with complete ORFs were retained for further analysis.

### Genome annotation

Repetitive elements were annotated using both *de novo* and homology-based approaches. RepeatMasker v2.1 ^68^ and the subrouting package RepeatProteinMask were used to identify known repeats based on Repbase. RepeatModeler v1.0.11 ^69^ (parameters: -species Haliotis -nolow -norna -no_is) and LTR_FINDER ^70^ generated a *de novo* repeat library, which was used by RepeatMasker v2.1 ^68^ to find additional repeats. Tandem repeats were identified by Tandem Repeat Finder v4.07 ^71^ with the following parameters: “Match = 2, Mismatch = 7, Delta = 7, PM = 80, PI = 10, Minscore = 50”.

For protein-coding genes, evidence from three approaches was integrated: (1) *de novo* prediction with AUGUSTES v3.2.1 ^72^ using the “seahare” model; (2) Homology-based prediction by aligning proteins from 8 molluscan genomes, *Aplysia californica* (GCA_000002075.2), *Biomphalaria glabrata* (GCA_947242115.1), *Crassostrea gigas* (GCA_011032805.1), *Lottia gigantea* (GCA_000327385.1), *Octopus bimaculoides* (GCA_001194135.2), *Pecten maximus* (GCA_902652985.1), *Pomacea canaliculate* (GCF_003073045.1), and *Pomacea maculate* (GCA_003073045.1), to the genome using tBLASTN v2.7.1 ^73^ (e-value 1e-5) and defining gene models with GeneWise ^74^; (3) RNA-seq support by aligning transcripts to the genome using BLAT v35 ^75^ and PASA ^76^ to generate gene models. The three lines of evidence were combined using EvidenceModeler v1.1.1 ^77^ to obtain a non-redundant gene set. Functional annotation was performed by scanning the gene set against databases including InterPro, GO, KEGG, UniProt, COG, Pfam, and NR using InterProScan v5.30-69.0 ^78^.

### Phylogenetics and demographic history

High-quality genomes of 14 mollusks were retrieved from NCBI, including *Haliotis rufescens* (GCA_023055435.1), *Haliotis laevigata* (GCA_008038995.1), *Haliotis rubra* (GCA_003918875.1), *Lanistes nyassanus* (GCA_004794575.1), *Marisa cornuarietis* (GCA_004794655.1), *Pomacea canaliculata* (GCF_003073045.1), *Pomacea maculate* (GCA_003073045.1), *Biomphalaria glabrata* (GCA_947242115.1), *Achatina immaculata* (GCA_009760885.1), *Achatina fulica* (http://gigadb.org/dataset/100647), *Cepaea nemoralis* (GCA_014155875.1), *Aplysia californica* (GCA_000002075.2), *Lottia gigantea* (GCA_000327385.1), and *Mizuhopecten yessoensis* (GCA_002113885.2). Orthologous genes were identified by performing ALL-to-ALL reciprocal BLAST searches using Diamond v2.0.4 ^79^. Individual gene trees and a supermatrix tree were constructed using IQ-TREE v1.6.12 ^80^ after alignment with MUSCLE v3.8.425 ^81^ and PAL2NAL v14 ^82^. A coalescent-based species tree was generated with ASTRAL v5.5.9 ^83^.

Divergence times were estimated using MCMCtree in PAML v4.9 ^84^, with calibrations from fossils and TimeTree ^85,86^: the most recent common ancestor (MRCA) of *Lanistes nyassanus* and *Pomacea canaliculata* >150Mya; MRCA of *Lottia gigantea* and *Aplysia californica*: 470.2-531.5 Ma; MRCA of *Haliotis iris* and *Haliotis rubra*: >84.9 Ma; MRCA of *Biomphalaria glabrata* and *Achatina fulica*: 177-187 Ma; MRCA of *Aplysia californica* and *Achatina immaculata*: 91-430 Ma; MRCA of *Pomacea canaliculata* and *Achatina immaculata*: 334-489 Ma; MRCA of *Haliotis iris* and *Achatina immaculata*: 401-507 Ma; MRCA of *Mizuhopecten yessoensis* and *Achatina immaculata*: 520-568 Ma.

The genome-wide substitution rates for each lineage were estimated using R8s v1.8.1 ^87^ with penalized likelihood and the same dataset and calibration points as above. For demographic history, SNPs were identified from aligned Illumina reads using SAMtools ^88^ (parameters: mpileup -q 20 -Q 20). Demographic changes were reconstructed using PSMC v0.6.5 ^89^ with a generation time of 4 years ^90^.

### Chromosome evolution analysis

Six gastropod species were selected: *Patella Vulgata* (GCF_932274485.2), *Gigantopelta aegis* (GCF_016097555.1), *Gibbula magus* (GCA_936450465.1), *Haliotis iris* (this study), *Achatina immaculata* (GCA_009760885.1), and *Pomacea canaliculate* (GCF_003073045.1). Syntenic blocks between chromosomes were identified using MCScanX v1.1 ^91^ (parameters: “-s 3 -w 25 -b 2 -m 100 -k 200 -g - 0.2”) given the low conservation between molluscan genomes. Ancestral karyotypes were constructed using ANGeS v1.01 ^92^ based on synteny to examine chromosomal evolution across lineages.

### Gene family analysis

Protein sequences from the 15 mollusks above were clustered using OrthoFinder v2.3.1 ^93^. Expansions and contractions of orthogroups were analyzed using CAFÉ v4.0.1 ^94^ with default parameters. Significance was determined based on both family-wide and Viterbi *P*-values < 0.01 ^95^.

### Identification of homeobox and melanogenesis genes

The protein sequences of 11 *Hox* genes and three *ParaHox* genes as reported by Wang et al. ^34^ were retrieved from MolluscDB (http://mgbase.qnlm.ac). These sequences were aligned to each genome using BLAT v.35 ^75^ to identify all candidate hits, retaining matches over 50% of the query length. Targeted sequences plus 10 kb up-and downstream were extracted and genes were precisely predicted using Genewise v2.4 ^74^. Redundant sequences were removed to obtain a non-redundant homeobox gene entry, which was verified by searching against the NR database.

Genomes from 10 representative metazoan genomes, *Homo sapiens* (GCA_000001405.29), *Leucoraja erinacea* (GCA_028641065.1), *Lethenteron reissneri* (GCA_015708825.1), *Ciona intestinalis* (GCA_009617815.1), *Branchiostoma lanceolatum* (GCA_927797965.1), *Acanthaster planci* (GCA_001949145.1), *Drosophila melanogaster* (GCA_000001215.4), *Caenorhabditis elegans* (GCA_000002985.3), *Mizuhopecten yessoensis* (GCA_002113885.2), and *Hydra vulgaris* (GCA_022113875.1), were combined with that of *H. iris* to generate a simplified metazoan phylogeny. The emergence and loss of melanogenic genes across this phylogeny were assessed using the same approach as described above. Key melanogenic genes were identified mainly based on the KEGG pathway ko04916.

### Topologically associating domain (TAD) analysis and loop calling

Hi-C sequencing data representing spatial interactions between loci were used to examine *pitx* region contacts. Resolution was first assessed using HiC-Pro v3.1.0 ^96^ with parameters “BIN_SIZE = 5000 10000 15000 20000 25000 30000 35000 40000 100000 200000” to evaluate contact matrices and determine the highest resolution where >80% of bins show >1000 contacts.

TADs were then identified from this optimal resolution matrix using “hicFindTADs” in HiCExplorer v1.5.12 ^97^, which measures separation between adjacent regions, with parameters “--minDepth 30000 --maxDepth 100000 --step 10000 --thresholdComparisons 0.05 --delta 0.05 --correctForMultipleTesting fdr”. TADs were defined as local minima in these TAD separation scores. TADs were visualized using pyGenomeTracks v3.7 ^98^.

Hic loop calling was conducted using “hicDetectLoops” in HiCExplorer v1.5.12 ^97^ with parameters: “--maxLoopDistance 2000000 --windowSize 10 --peakWidth 6 -- pValuePreselection 0.05 --pValue 0.05”.

### Transcriptome differential expression analysis

RNA-seq reads from each *H. iris* tissue were filtered using fastp v0.23.1 ^59^ (parameters: -n 0), then aligned to the genome using Hisat2 v2.1.0 ^99^. Stringtie v2.2.1 ^100^ was used to calculate the reads count of each gene and quantify gene expression as TPM values based on the gene annotations GFF. Differentially expressed genes were identified using DESeq2 ^101^, requiring |log_2_FC| ≥ 1 and FDR < 0.05. GO enrichment analysis was performed to examine functions of differentially expressed genes (Chi.FDR < 0.05).

To identify the candidate differentially expressed non-coding elements in left and right mantles, transcripts obtained from mantles without ORFs were aligned to the genome using PASA ^76^ to define structures. These were filtered against protein-coding annotations, resulting in 141,450 non-coding elements active in the mantle (Supplementary Table S4). The paired-end oligo-dT libraries were used to capture the strand information (Supplementary Table S1). Finally, the annotation file of these non-coding elements (GFF) was used to quantify and identify their differentially expressed genes as described above for coding genes.

### DNA methylation analysis

Paired-end whole-genome bisulfite sequencing reads were filtered using fastp v0.23.1 ^59^ with default parameters. The *H. iris* genome was converted using “bismark_genome_preparation” in Bismark v0.23.0 ^102^, which performs in silico C->T and G->A conversions. Filtered reads were mapped to the converted genome using Bowtie2 v2.3.4.3 ^103^. Methylation information at all C positions was extracted using “bismark_methylation_extractor” in Bismark v0.23.0 ^102^ to obtain data for CpG/CHG/CHH contexts. Differentially methylated positions were identified using the R package “*methylKit*” ^104^ requiring at least 20% methylation change and FDR < 0.05.

### Quantitative analysis of genes

The mRNA expression of genes, *viz. pitx*, *lncRNA1*, *lncRNA2*, *tyr-1, tyr-2*, and *mitf*, was quantified by quantitative reverse transcription PCR (qRT-PCR) using a thermal cycler (Applied Biosystems, USA) and SYBR Green 2X Master Mix (Takara). Genes’ mRNA were amplified from abalone cDNA using specific primers (Supplementary Table S21). One of the genes, *GAPDH*, *EF-1a*, and *β-actin*, served as an internal control. PCR cycling conditions were: initial denaturation at 95°C for 30 s, followed by 40 cycles of 95°C for 5 s and 60°C for 30 s. Melting curves were analyzed after each run to ensure specificity. The 2^-ΔΔCT^ method was used to calculate the relative expression. One-way ANOVA with unpaired two-tailed *t*-tests was used for statistical analysis.

### RNA interference of lncRNAs and *mitf*

To investigate the effects of lncRNAs and *mitf* knockdown, gene-specific small interfering RNAs (siRNAs) and corresponding negative control (NC) siRNAs were designed and synthesized by commercial providers. For RNA interference (RNAi) experiments, rainbow abalones were allocated to RNAi and NC groups (typically *n* = 6-9 per group). For lncRNAs, each abalone received an initial injection of siRNA (50 µg per animal) dissolved in PBS, followed by a second identical injection 18 hours later to enhance knockdown efficacy. Mantle tissues were collected 12 hours post-second injection. For *mitf*, each abalone received an initial injection of siRNA (30 µg per animal) dissolved in PBS, followed by a second identical injection 12 hours later to enhance knockdown efficacy. Mantle tissues were collected 24 hours post-second injection. The knockdown efficiency of the target lncRNAs or *mitf*, and the relative expression levels of their respective downstream genes (*pitx* for lncRNA interference; *tyr-1* and *tyr-2* for *mitf* interference), were quantified using qRT-PCR. Statistical significance of differences between RNAi and NC groups was determined using independent samples t-tests. All siRNA and qRT-PCR primer sequences are detailed in the Supplementary Table S21.

### Histological staining and *in situ* hybridization

For the histological survey, we employed conventional paraffin sectioning techniques to prepare histological sections of the left anterior mantle lobe in the rainbow abalone. The preparation of sections and hematoxylin-eosin (HE) staining followed previously published protocols ^105^ and were meticulously optimized based on the preservation conditions of the samples (e.g., sample fixation, section thickness, etc.).

Fluorescent *in situ* hybridization was performed and optimized according to the method described by Kishi et al. ^106^. Briefly, after rehydration, digestion, and pre-hybridization (37℃, 1h), left mantle tissue sections were hybridized with hybridization probes (500nM, designed and synthesized by commercial providers; Supplementary Table S11) at 40℃ overnight. And then the probes were washed away with pre-warmed 2×SSC, 1×SSC, and 0.5×SSC. After extensive washing, sections were added branch hybridization solution and held at 40°C for 45min in a humid chamber. The sections were then washed with 2×SSC and rinsed in 1×PBS. Subsequently, the sections were incubated in fluorescent hybridization solution (1:400 iF488-Tyramide or Cy3 dye) and incubated at 42°C for 3 hours. After washing consecutively with 1×PBS, the sections were counterstained with DAPI reagent. Finally, sections were visualized under a fluorescence microscope (NIKON, ECLIPSE CI), and images were captured. Fluorescence signals were acquired using appropriate filter configurations: blue fluorescence (typically DAPI for nuclear counterstain, UV excitation 330-380nm, emission ∼420nm), green fluorescence (generated via iF488-Tyramide signal amplification, excitation ∼470-495nm, emission ∼510-530nm), and red fluorescence (CY3 conjugated probe, excitation ∼510-560nm, emission ∼590nm).

Whole-mount *in situ* hybridization (WISH): The probes of *lncRNA1* and *pitx* were synthesized using specified primers (Supplementary Table S21) from the cDNA library of the left lobe of the mantle. Plasmid-derived riboprobes were synthesized using T7 polymerases, tagged with digoxigenin labels ^107^. Concurrently, abalone embryos at various developmental stages, including artificial fertilization, 2-cell stage, 4-cell stage, 16-cell stage, morula, gastrula, non-hatched trochophore, trochophore, early veliger, and late veliger stages, were sequentially collected and rinsed with RNase-free water. The embryos underwent thorough washes with a dedicated solution and DEPC-treated water, followed by Proteinase K digestion. Subsequently, they were washed with 0.1 mol/L glycine solution, PBS, and 5×SSC solution. The hybridization and immunological detection procedures were carried out as previously described ^34,108^.

### *lncRNA1*-miRNAs interactions

To identify miRNAs interacting with the target *lncRNA1* in abalone, HyPro-proximity miRNA pulldown ^38^ was performed by Aksomics (Shanghai, China). Briefly, tissues were ground in liquid nitrogen, and a portion was reserved as the input control. For the experimental group, samples underwent DSP crosslinking, ethanol permeabilization, and hybridization with a custom DIG-conjugated DNA probe specific to the target *lncRNA1* (Supplementary Table S11). A control HyPro pulldown was performed in parallel, purportedly using random sequence DNA probes within the HyPro workflow. The HyPro fusion enzyme (APEX2-DIG binding protein) was then incubated, followed by proximity-dependent biotinylation of adjacent RNA molecules using biotin-phenol and H₂O₂. After cell lysis, total RNA was extracted, and biotinylated RNAs were captured using streptavidin magnetic beads, followed by RNA elution and purification. Small RNA libraries were constructed from the enriched RNA fractions and the control RNA using the NEB Multiplex Small RNA Library Prep Set for Illumina (New England Biolabs, USA), involving 3’ and 5’ adapter ligation, cDNA synthesis, PCR amplification, and size selection of 135-155 bp PCR products (corresponding to 15-35 nt small RNAs). Library quality was assessed using an Agilent 2100 Bioanalyzer. Sequencing was performed on an Illumina platform using the TruSeq Rapid SR Cluster Kit. Raw sequencing reads were processed with Cutadapt ^109^ to remove adapters, and tags ≥17 nt were retained as trimmed reads for downstream analysis.

Following quality control, the trimmed reads were mapped to the *H. iris* genome using Bowtie2 ^103^ with the “--very-sensitive-local” preset parameter. Alignments with a mapping quality score greater than 20 were retained for further analysis. The SAMtools ^88^ and Bedtools ^110^ software suites were subsequently employed to derive genomic coordinates and quantify read counts for the successfully mapped reads. For comparative analysis, read counts corresponding to identified miRNAs in both the experimental and control groups were normalized to Counts Per Million (CPM) mapped reads. To aid in the assessment of differential miRNA enrichment, a fold difference (FD) index was calculated using the formula: FD = log₁₀(|CPM_IP – CPM_nc|) × log₂(CPM_IP/ (CPM_nc + 1)), where CPM_IP denotes the CPM value for the experimental sample and CPM_nc denotes the CPM value for the negative control sample.

### Dual-luciferase reporter assays

Due to the lack of molluscan cell lines for transfection, human HEK293T cells were used for heterologous expression of abalone proteins. Cells were cultured in high-glucose Dulbecco’s modified Eagle’s medium (HyClone, USA) supplemented with 10% FBS (Gibco, USA) and 1x penicillin/streptomycin (Solarbio, China) at 37°C with 5% CO_2_ and passaged every 2-3 days.

To investigate the direct interaction between miRNAs and their putative targets, reporter constructs were generated. Specifically, predicted target sequences within the abalone *pitx* mRNA and *lncRNA1* were commercially synthesized (Sangon Biotech, China; Supplementary Table S21). These synthetic DNA fragments were subsequently cloned into the pmir-GLO Dual-Luciferase miRNA Target Expression Vector (Promega, USA) via *Xho*I/*Sal*I restriction sites, yielding the *pitx* and *lncRNA1* mRNA reporter vectors, respectively. Chemically synthesized miRNA mimics for *miRNA1*, *miRNA2*, *miRNA3*, and a negative control (NC) mimic (Supplementary Table S21) were procured from GenePharma (Shanghai, China). For target verification, HEK293 cells were co-transfected with either the *pitx* mRNA reporter vector or the *lncRNA1* reporter vector, along with the respective miRNA mimics (*miRNA1*, *miRNA2*, or *miRNA3*) or the NC mimic. The pRL-CMV Renilla luciferase plasmid (Promega) was included in all co-transfections as an internal reference. Luciferase activities were measured using the Dual-Luciferase® Reporter Assay System (Promega), and firefly luciferase activity was normalized to Renilla luciferase activity. Data from miRNA mimic-transfected groups were compared against the NC group using an unpaired two-tailed Student’s t-test.

To evaluate the regulatory activity of abalone MITF on target gene promoters, reporter and expression plasmids were constructed. Promoter regions of rainbow abalone *tyr-1* or *tyr-2* genes were amplified by PCR from cDNA using KOD One™ PCR Master Mix (TOYOBO, Osaka, Japan) with specific primers (Supplementary Table S21) designed based on gene sequences and predicted protein domains. Purified PCR products (DNA Gel Extraction Kit, Sangon Biotech) were inserted into the linearized pGL6-TA reporter plasmid (Beyotime, Shanghai, China) *via* In-Fusion cloning. Concurrently, an expression plasmid encoding abalone MITF was generated by cloning the *mitf* coding sequence into the pCMV-Myc expression vector (Clontech, Mountain View, CA, USA) using In-Fusion cloning. For reporter assays, HEK293T cells, seeded in 24-well plates, were transfected at 60-70% confluency. Transfections included the appropriate promoter-reporter plasmid, the pRL-TK Renilla luciferase internal control plasmid (Promega), and either the MITF expression plasmid or an empty pCMV-Myc vector as a control, using Lipofectamine 3000 (Thermo Fisher Scientific, Waltham, MA, USA). Cells were harvested 24-36 hours post-transfection. Firefly and Renilla luciferase activities were quantified using the Dual-Glo® Luciferase Assay System (Promega). Firefly luciferase activity was normalized to Renilla luciferase activity and expressed as fold induction relative to cells transfected with the empty vector control. All promoter activity experiments were performed in triplicate.

### Abalone embryonic transcriptome

Due to the inaccessibility of *H. iris* embryos, *Haliotis discus* embryos representing 14 developmental stages were used as a substitute: artificial fertilization, 2-cell, 4-cell, 8-cell, 16-cell, morula, gastrula, non-hatched trochophore, trochophore, early veliger, late veliger, peristomial shell larva, differentiation epipodes, and emergence of the 1st breathing pore. RNA was extracted from whole flash-frozen embryos for RNA-seq. Data was analyzed similarly to the tissue transcriptomes above.

### Synteny analysis of *pitx* and *lncRNA1*

To investigate the conservation and structural context of *pitx* and *lncRNA1*, RNA-seq reads for species pertinent to the chromosome evolution analysis (Supplementary Table S22) were retrieved from the NCBI database. Raw reads were subjected to quality filtering using fastp v0.23.1 ^59^ with the parameter “-n 0”. The filtered reads were then mapped to their respective reference genomes using HISAT2 v2.1.0 ^99^. Initial expression levels of the *pitx* gene were assessed using StringTie v2.2.1 ^100^. For RNA-seq reads exhibiting *pitx* expression, they were further processed with StringTie v2.2.1 ^100^ for reference-based transcript assembly. The resulting transcripts were subsequently refined using PASA ^76^ to identify accurate gene structures. Finally, the syntenic relationship between *pitx* and *lncRNA1* across these species was visualized using the IGV browser, based on the gene structure files (GFF, from PASA) and alignment files (bam, from HISAT2) generated from the aforementioned analyses.

### Yeast one-hybrid assays

To validate *mitf* regulation of *tyr* genes, yeast one-hybrid assays were performed. Promoter regions of *tyr-1* and *tyr-2* containing predicted MITF binding sites were PCR amplified (Supplementary Table S21) and cloned into the pLacZi vector. *mitf* coding sequences were cloned into the pB42AD vector (Supplementary Table S21). Plasmids were co-transformed into the yeast strain EGY48 and transformants selected on SD/-Trp/-Ura plates. Positive colonies were screened for reporter activation on chromogenic medium with X-gal.

### Transmission electron microscopy

To observe melanin particles in the abalone mantle, the ultrastructure was examined by transmission electron microscopy. Mantle tissue pieces (< 1 mm^3^) from fresh abalone were fixed in 2.5% glutaraldehyde. After rinsing with 0.1M phosphate buffer, gradient acetone dehydration was performed. Tissue pieces were resin-embedded and cured at 37°C, 45°C, and 60°C sequentially. Ultrathin 70 nm sections were cut using a Reichert-Jung ultracut E ultramicrotome (Austria) and collected on copper mesh. Sections were stained with uranium acetate and lead citrate before imaging on a JEM1200 electron microscope.

## Acknowledgments

The authors thank our collaborators, Xiangxiang Wang, Mingliang Hu, Hui Shi, Yanjie Zhang, Liyan Qiao, Hongce Song, Jilv Ma, Xi Chen and Yanfei Ma, for their advice and discussions during this project.

## Funding

This research was supported by the Key Research and Development Program of Shandong Province (2022LZGC015); the National Key R&D Program of China (2022YFC3400300), by the Fundamental Research Funds for the Central Universities, Northwestern Polytechnical University (D5000220464), Special Funds for the Taishan Scholar Project of Shandong Province, China (tstp20240518); Modern Agricultural Industry Technology System of Shandong Province, China (SDAIT-14-01), and the National Natural Science Foundation of China (32370452).

## Author Contributions

X.W., C. F., Q.Q., and K.W. conceived and led the study; C. F., X.W., X.G., and K.W. designed this project and research aspects; X.W. and M.Z. performed sample collection; C.L., P.X., S.L., C.Z., W.Xu, and C.F. performed data analyses, including sequencing data preparation, genome assembly, annotation, and subsequent comparative genomics analyses; P.X., M.Z., B.H., T.C., Y.H., L.W., X.Z., Y.L., W.Xue, Z.Y., W.Y., P.Z., L.L., M.Y., D.T., and J.Q. carried out experimental analysis and discussion, including histoembryology, molecular biology, and cell biology. C.F., X.W., C.L., P.X., M.Z., B.H., T.C., K.W., and X.G. wrote the manuscript. All authors were involved in the revision of the manuscript and gave the final approval for publication.

## Competing Interests

The authors declare no competing interests.

## Data and Materials Availability

Genome assembly and raw sequencing data have been deposited in the CNGB Sequence Archive (CNSA) of China National GeneBank Database (CNGBdb) with accession number (CNP0004800). All other study data are included in the article and/or supporting information.

## Notes

### Competing Interest Statement

The authors have declared no competing interest.

